# An extreme mutational hotspot in *nlpD* depends on transcriptional induction of *rpoS*

**DOI:** 10.1101/2024.07.11.603030

**Authors:** Andrew D. Farr, Christina Vasileiou, Peter A. Lind, Paul B. Rainey

**Author notes:** Corresponding authour.

## Abstract

Mutation rate varies within and between genomes. Within genomes, tracts of nucleotides, including short sequence repeats and palindromes, can cause localised elevation of mutation rate. Additional mechanisms remain poorly understood. Here we report an instance of extreme mutational bias in *Pseudomonas fluorescens* SBW25 associated with a single base-pair change in *nlpD.* These mutants frequently evolve in static microcosms, and have a cell-chaining (CC) phenotype. Analysis of 153 replicate populations revealed 137 independent instances of a C565T loss-of-function mutation at codon 189 (CAG to TAG (Q189*)). Fitness measures of alternative *nlpD* mutants showed molecular parallelism to be unconnected to selective advantage. Recognising that transcription can be mutagenic, and that codon 189 overlaps with a predicted promoter (*rpoSp*) for the adjacent stationary phase sigma factor, *rpoS*, transcription across this promoter region was measured. This confirmed *rpoSp* is induced in stationary phase and that C565T mutation caused significant elevation of transcription. The latter provided opportunity to determine the C565T mutation rate using a reporter-gene fused to *rpoSp*. Fluctuation assays demonstrate the C565T mutation rate to be 5,700-fold higher than expected. In *Pseudomonas*, transcription of *rpoS* requires the positive activator PsrA, which we show also holds for SBW25. Fluctuation assays performed in a Δ*psrA* background showed a 60-fold reduction in mutation rate confirming that the elevated rate of mutation at C565T mutation rate is dependent on induction of transcription. This hotspot suggests a generalisable phenomenon where the induction of transcription causes elevated mutation rates within defining regions of promoters.

## Introduction

Mutation fuels evolution by natural selection. Despite the long-term benefits of mutation for life on earth, most mutations are neutral or deleterious (Eyre-Walker & Keightley, 2007), and the rate at which they occur is tightly regulated (Lynch, 2010; Sniegowski, Gerrish, Johnson, & Shaver, 2000). In bacteria, point mutations to any one base-pair occur approximately once every 1 × 10^10^ cell divisions, with the lowest average mutation rates occurring in *Pseudomonas fluorescens* SBW25 (Long, Miller, Williams, & Lynch, 2018). However the rate of mutation varies significantly between regions of genomes (extensively reviewed (Horton & Taylor, 2023)), with some nucleotide positions – referred to as “mutational hotspots” – subject to high frequency change.

The mechanistic causes of mutational hotspots are often idiosyncratic and unpredictable. Simple sequence repeats can undergo slipped-strand mis-repairing (Levinson & Gutman, 1987), resulting in high-frequency gain or loss of repeats (Moxon, Rainey, Nowak, & Lenski, 1994). Imperfect inverted repeat sequences (‘quasi-palindromes’) may induce template-switching, with a range of mutational effects (Lovett, 2017). Other hotspots emerge from interactions with transcriptional machinery. In a study of mutation rates at *thyP3* in *Bacillus subtilis*, Sankar and colleagues measured high levels of thymine to cytosine point mutations (at a rate of ∼1 × 10^−8^ per cell per generation) at a single base pair directly upstream of the transcription start site (Sankar, Wastuwidyaningtyas, Dong, Lewis, & Wang, 2016). This was enhanced by the activity of the promoter and orientation of the gene, leading to the suggestion that stalled RNA polymerase can allow deamidation of nucleotides on template strands. Such specific mutagenic processes can interact with regional mutational biases, such as genomic strandedness and location, to enhance mutation rates (Horton, Flanagan, Jackson, Priest, & Taylor, 2021; Shepherd, Horton, & Taylor, 2022), but also any process that biases the spectrum of mutations in the genome (Gifford et al., 2024; Krasovec et al., 2018; Sane, Diwan, Bhat, Wahl, & Agashe, 2023). The multitude of mechanistic causes of hotspots has a singular consequence: they increase the production of variants presented to selection (McCandlish & Stoltzfus, 2014), often explaining instances of parallel evolution (Lind, Libby, Herzog, & Rainey, 2019).

A remarkable example of parallel evolution was previously identified across multiple independent cultures and during independent studies using the model organism *Pseudomonas fluorescens* SBW25 (Gallie et al., 2019; Lind, Farr, & Rainey, 2017). Parallelism was evident from repeated observation of identical cytosine (C) to thymine (T) mutations at base pair 565 (C565T) of *nlpD*. The mutation alters codon 189 (sequence CAG – encoding glutamine) and results in a premature stop codon (sequence TAG). These mutants – arising in statically incubated microcosms of KB media due to a measurable fitness advantage over ancestral genotypes (Lind et al., 2017) – were identified by subtle changes to colony morphology and a cell-chaining (CC) phenotype. The CC phenotype is readily explained by the predicted function of NlpD *–* a protease that enables activation of amidases that cleave peptidoglycan during cellular division (Yakhnina, McManus, & Bernhardt, 2015). Depending on the presence of related genes, loss-of-function mutations in NlpD can result in incomplete cell separation in genera such as *Escherichia* (Tsang, Yakhnina, & Bernhardt, 2017) and *Pseudomonas* (Yakhnina et al., 2015). It was previously suggested that repeated detection of the C565T mutation in *nlpD* might be explained by a secondary function encoded by *nlpD* (Lind et al., 2017), namely, that nested within *nlpD* is the primary promoter for the downstream gene, *rpoS* (we refer to the promoter of *rpoS* as ‘*rpoSp’*) (Fujita, Tanaka, Takahashi, & Amemura, 1994; Kojic, Aguilar, & Venturi, 2002; Venturi, 2003). Assuming this mutation also alters transcription of *rpoS*, the combination of NlpD truncation and altered RpoS expression might explain the post-selection observation of mutations in *nlpD* (Lind et al., 2017).

In this study, we build upon previous work and demonstrate that a single codon in *nlpD* is prone to mutate at high frequency. Assessment of the fitness of additional *nlpD* mutants shows that molecular parallelism at C565T is not caused by selection. Characterisation of the *rpoS* promoter led to construction of a mutational reporter to measure mutation rate at C565T with further work demonstrating that elevated mutation is transcription-dependent.

## Results

### Extreme molecular parallelism within *nlpD*

We previously reported mutational parallelism of the C565T mutation following adaptive radiation of the SBW25 Δ*wss* strain in statically incubated microcosms (Lind et al., 2017). We wanted to understand the mechanistic basis of this molecular parallelism. However, we first repeated the original experiment but with greater replication: 153 static broth microcosms were inoculated with SBW25 Δ*wss* and screened for the presence of CC mutants. SBW25 Δ*wss* was used to limit the evolution of ‘Wrinkly Spreader’ morphotypes that readily evolve in static microcosms inoculated with SBW25 and otherwise obscure detection of CC types (Rainey & Travisano, 1998). The cultures were diluted, spread on agar plates, and subsequent colonies were screened for the aberrant cellular chaining (CC) morphology characteristic of *nlpD* mutants. From 153 populations, CC mutants were detected in 139 populations, with sequencing showing that 137 contained the C565T mutation (Fig. 1A), one a G539A mutation (resulting in a G180D non-synonymous mutation) and one a frame shift (fs) insertion inside a homopolymeric tract (base pairs 705-708; Q237fs). While additional mutations to *nlpD* can arise in static microcosms, the predominant cause of the CC type was the C565T mutation, produced by an extreme degree of molecular parallelism.

**Figure 1:**
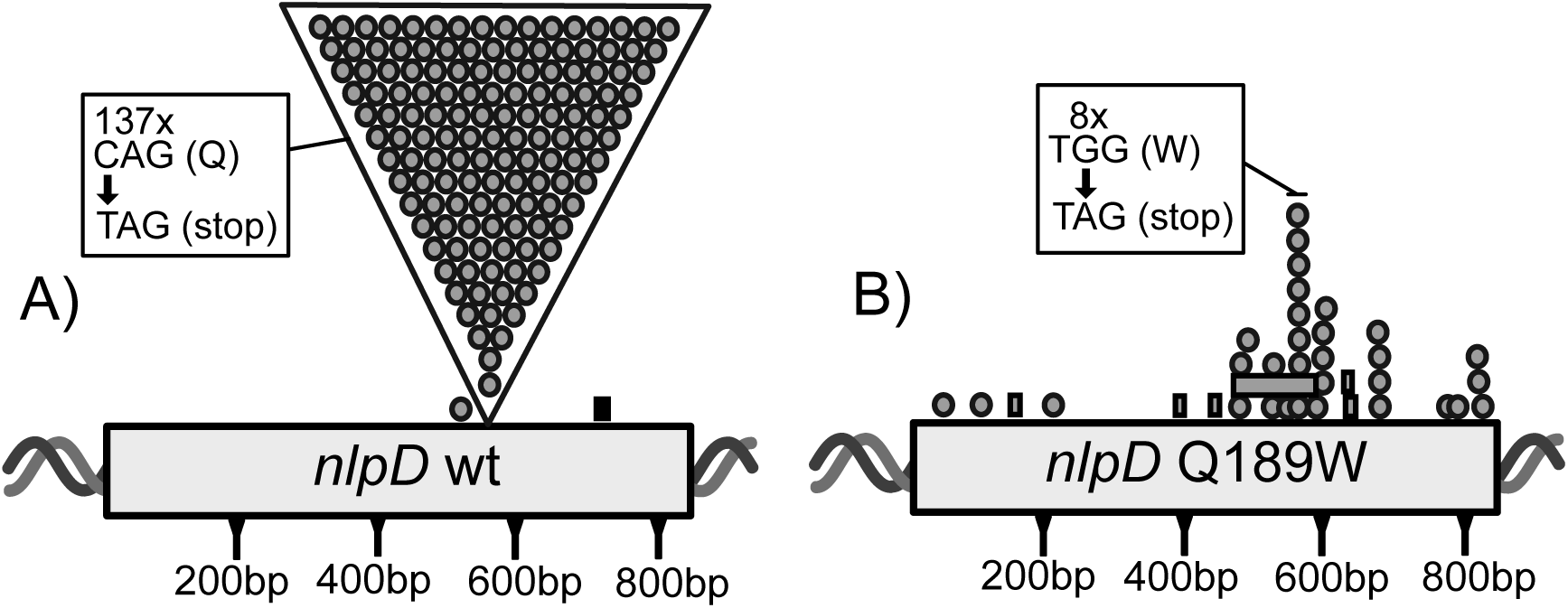
The position of CC mutations found within *nlpD*. **A) Distribution of *nlpD* mutants evolved from SBW25 Δ*wss*.** Depicted is the position of *nlpD* mutants evolved from SBW25 Δ*wss* cultured in static microcosms. The majority (137) of mutants were identical C to T mutations (gray circles in large triangle) at base pair 565 of *nlpD* (C565T). The C565T mutation results in a CAG to TAG sequence change at codon 189 (Q189*; see box). Two alternative nonsynonymous mutations were identified, a point mutation and an insertion (black rectangle). **B) Distribution of *nlpD* mutants evolved from SBW25**Δ***wss nlpD* Q189W.** This distribution of mutations changes when the founding genotype is SBW25 Δ*wss nlpD* C565T A566G (Q189W), with the sequence of codon 189 being TGG. Evolution from this genotype resulted in point mutations (grey circles) and deletion mutations (grey rectangles) spread across the open reading frame of *nlpD*. A degree of molecular parallelism remains, with eight mutants having the G566A mutation (resulting in a TGG to TAG sequence change at codon 189; see box).

A previous study (Gallie et al., 2019) had serendipitously obtained a *nlpD* mutant that firstly acquired the C565T mutation, followed by a second mutation at nucleotide 566 (A566G) delivering a CAG to TGG sequence change at codon 189 of *nlpD*. This codon change resulted in a glutamine to tryptophan substitution – Q189W – at codon 189. Given that the TGG mutation at codon 189 was not CC (Gallie et al., 2019), we asked whether CC mutants might still arise by disproportionate changes at codon 189 caused specifically by reverting G566A mutations (resulting in a sequence change of TGG back to TAG). To this end, codon 189 of *nlpD* in SBW25 Δ*wss* was altered from CAG to TGG and the resulting mutant propagated in 54 static broth microcosms. CC mutants were detected in 38 microcosms, with 37 harbouring mutations in *nlpD*. Of these, eight acquired the reverting G566A mutation (resulting in TAG stops at codon 189), with the remaining mutations scattered throughout the gene (Fig. 1B and Table S1). This shows that a subtle change in the sequence of codon 189 dramatically reduces (but does not abolish) the extreme parallelism previously observed.

### Fitness of C565T mutants does not explain parallel mutation

Armed with a suite of additional CC mutants carrying defects in *nlpD* other than C565T, it was possible to ask whether the C565T mutation conferred a disproportionate selective advantage that might explain its repeated occurrence. The fitness of six early termination mutations derived from the Q189W (TGG) mutant (Table S1), as well as the Q237fs mutant evolved from SBW25 Δ*wss*, were compared to the C565T mutant. For genotypes derived from Q189W, codon 189 was restored to the ancestral CAG sequence, to eliminate any effects of the Q189W mutation. Five of the most 3’ *nlpD* mutations all had significantly lowered fitness compared to the control competitions of C565T against itself (Fig. 2; p<0.0001; Welch’s one-way ANOVA, F(8,25.652) = 58.649, p<0.0001, pairwise difference assessed with Tukey HSD). The two 5’ *nlpD* mutations (stop and frameshift mutations at codons 17 and 132) were not significantly different from the control. The observation of alternative *nlpD* mutants expressing equivalent fitness to the C565T mutant suggests that while many mutations of *nlpD* have lower fitness compared to C565T mutants, the repeated evolution of the C565T mutant is not explained by a competitive advantage attributable to the C565T mutation.

**Figure 2:**
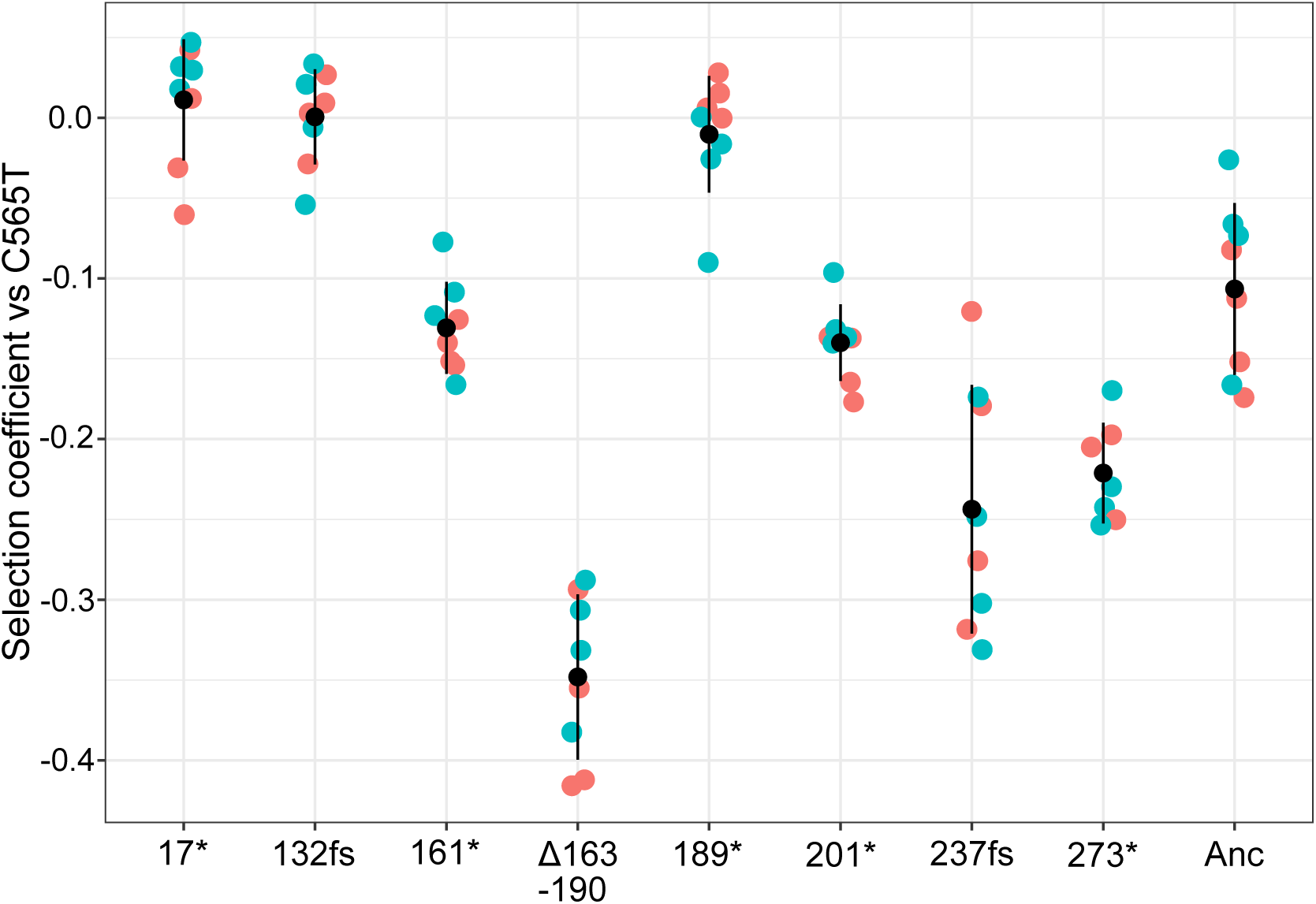
Fitness of CC mutants containing mutations in *nlpD* relative to the C565T mutant. Each *nlpD* mutation (labelled here with amino acid alteration) was reconstructed into a fluorescently labelled *SBW25* Δ*wss* background and was competed against the C565T mutant (with initial ratio 1:1). A genotype with ancestral *nlpD* (anc) was also included, showing a mild fitness benefit of the C565T mutant compared to the ancestor. Reciprocal pairwise competitions were used, with the indicated mutation in genotypes expressing either *gfp* (green) or *mScarlet* (red) florescence. Each competition involved 8 replicates, black dots represent the mean values and error bars are one standard deviation from the mean.

### Transcription from the *rpoS* promoter (*rpoSp*) is elevated in stationary phase and is strengthened by the C565T mutation

An alternative explanation for the near-deterministic evolution of the C565T mutants is that the region around codon 189 of *nlpD* mutates at a high rate, resulting in the C565T mutation being presented to selection at a high frequency. Could the presence of *rpoSp* – inferred as proximal to the mutation from promoter mapping in other pseudomonads (Fujita et al., 1994; Kojic et al., 2002) – define a mutational hotspot? Confirmation and characterisation of this promoter could also prove useful: any potential alteration of transcription by the C565T mutation may allow development of a mutation detection system and measurement of mutation rates.

Levels of transcript downstream from *rpoSp* were measured by RT-qPCR, using transcript directly upstream of *rpoSp* as a reference (Fig. 3A). These measures were performed using stationary phase cultures of WT *nlpD*, the C565T mutation and other *nlpD* loss-of-function mutations reconstructed in SBW25 Δ*wss.* These additional mutations were included to assess whether the CC phenotype might result in alteration of transcription from *rpoSp*. Shaken cultures were used as opposed to static cultures to homogenously induce stationary phase. These measures revealed a ∼7-fold increase in transcript produced across *nlpD* (Fig. 3B), indicating the presence and expression of *rpoSp* in stationary phase cultures. Remarkably, while three mutations distal to *nlpD* had similar levels of transcript to the ancestral SBW25 Δ*wss*, the C565T mutation significantly increased the relative level of transcription across *rpoSp* by a further ∼6.6-fold ((Fig. 3B); p<0.0001; one-way ANOVA, F4,15=109.5, p<0.0001, pairwise difference assessed with Tukey HSD). The observation that the C565T mutation results in a dramatic increase in transcription from *rpoSp* allowed construction of a mutational reporter, such that the mutation rate of the C565T mutation could be measured without the confounding influence of changes to NlpD.

**Figure 3:**
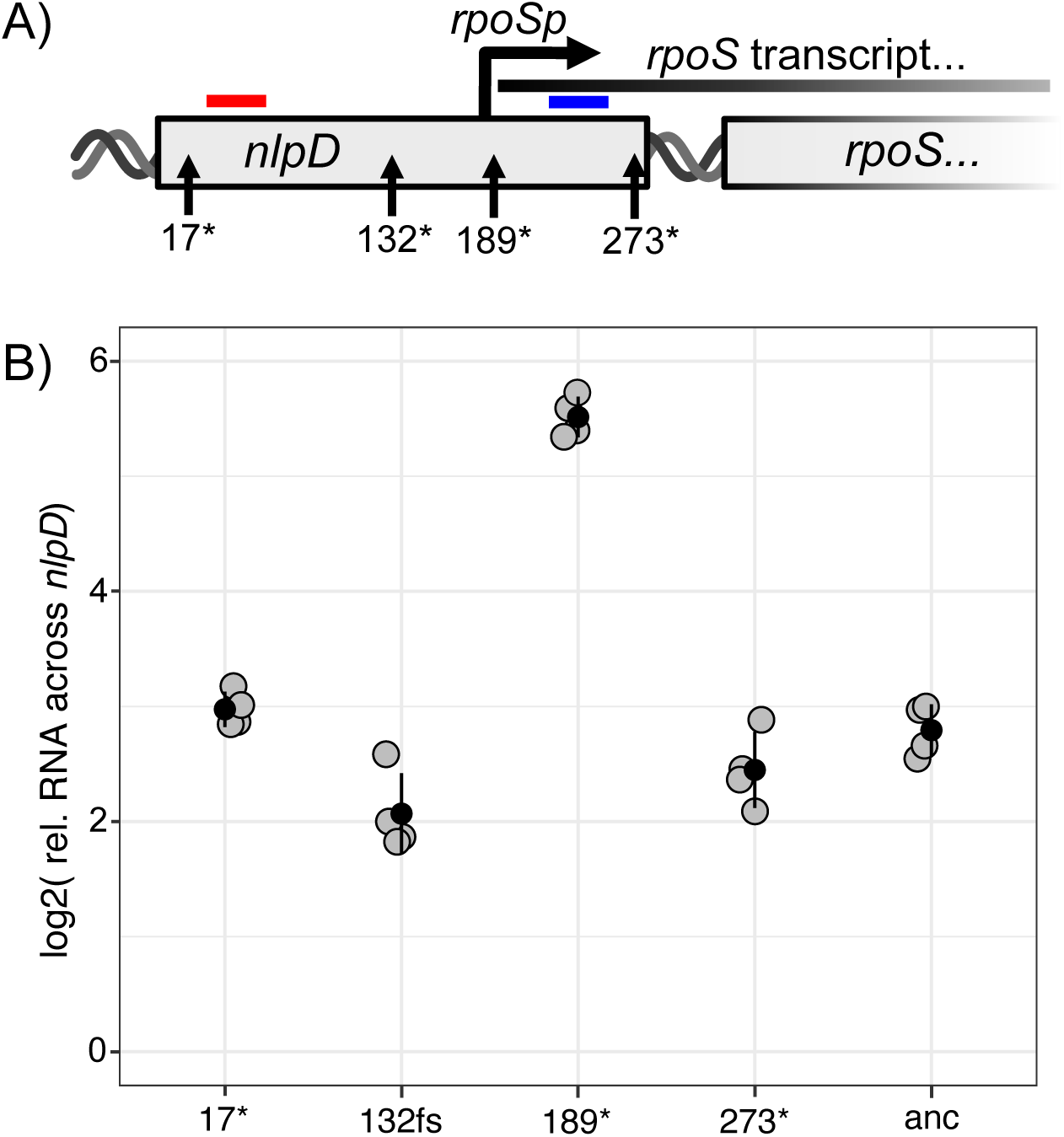
The Q189 mutation elevates transcription from *rpoSp* compared to other *nlpD* mutants or the ancestral sequence. **A) The location of the four mutants, the regions with quantified transcript, and the predicted region of *rpoSp*. Transcript levels were measured downstream (blue bar) relative to upstream (red bar) of *rpoSp*.** Transcription across *nlpD* was measured for mutants with loss-of-function mutations at codon 17 (TGA), 132 (a frameshift), 189 (TAG) or 273 (TAA), as well as the ancestral (anc) SBW25 *Δwss* **B) Measures of transcript across *nlpD* for the five *nlpD* genotypes.** Strains were grown to stationary phase, mRNA was extracted and reverse transcribed and qPCR performed on regions up and down stream of *rpoSp*. The plot depicts the relative amount of transcript downstream of *rpoSp* relative to upstream. The Q189* mutation causes a large (∼42 fold) increase in transcript from *rpoSp*. Black dots represent the mean values of four biological replicates (gray dots) and error bars one standard deviation from the mean.

### The C565T mutation rate is orders of magnitude higher than the average C to T mutation rate

We designed and constructed a mutational reporter that would allow measurement of the frequency of C565T mutants in growing populations of *P. fluorescens* SBW25. The construct consisted of a 401 bp portion of *nlpD* (with C565T central to this construct) fused to a kanamycin resistance gene. Upon a C565T mutation, the transcription from *rpoSp* is expected to increase, causing cells to become resistant to kanamycin and allowing identification of rare C565T mutations by colony growth on agar plates supplemented with kanamycin. The construct was integrated into the genome of SBW25 at a site distal to *nlpD* thus allowing identification of C565T mutations without interfering with *nlpD* function.

Fluctuation assays were then performed (Luria & Delbruck, 1943) on cultures of SBW25 harbouring the ‘*rpoSp-kan’* selectable reporter. Twelve independent cultures were grown from a small inoculum (∼1000 CFU mL^−1^) in shaken media for 22 h of growth until SBW25 cells had induced transcription from *rpoSp* (as confirmed by qPCR, see Fig. S1). Under these conditions, the mean frequency of the C565T mutant in this mutational reporter is once in ∼2 × 10^5^ cells in the background of SBW25 (Fig. 4A (left)). The mutation rate of the C565T mutation was 4.2 × 10^−7^ per base-pair per replication (95% CI = 4.5 – 3.9 × 10^−7^), as derived by a MSS Maximum Likelihood Method. Compared to previous measures of the average rate of C to T mutations in SBW25 (Long et al., 2018), this represents a ∼5700-fold increase in the rate of C to T mutations at base pair 565. Interestingly, these fluctuation assays revealed a mutational hotspot slightly broader than base-pair 565 of *nlpD*. Sanger sequencing of the mutant colonies identified 6 of 96 sequenced kanamycin-resistant colonies of SBW25 *rpoSp-kan* having a A564G mutation in the reporter. Presumably this mutation occurs in *nlpD* in populations of SBW25, but is not selected in static microcosms because it results in a synonymous mutation. Additional to fluctuation assays, we performed fitness assays and measured a near-neutral effect of these C565T mutation in the *rpoSp-kan* construct in growing cultures (Fig. S2). These measures suggest C565T mutants derived during fluctuation assays resulted solely from high mutation rates. This high rate of mutation to *rpoSp* will likely result in a high rate of mutation when *rpoSp* is nested within *nlpD*, which in the context of *nlpD* create CC mutants in wild populations of SBW25.

**Figure 4:**
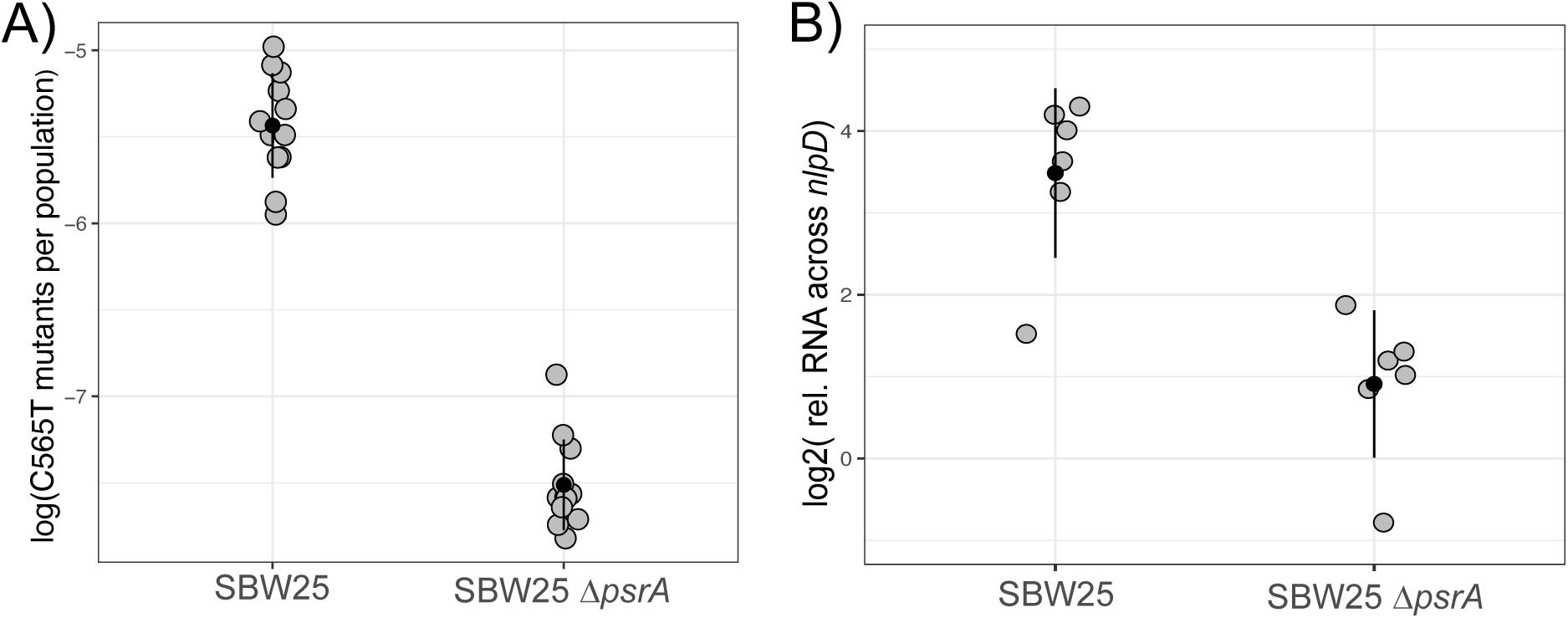
The frequency of the C565T mutation is associated with induction of *rpoSp*. **A) The frequency of C565T mutants identified using the *rpoSp-kan* reporter in populations of SBW25 and SBW25 Δ*psrA*.** SBW25 and SBW25 Δ*psrA* (both with the *rpoSp-kan* reporter were grown from a small inoculum (∼1000 CFU mL^−1^) for 22 h and samples plated on agar with and without selective concentration of kanamycin to measure the frequency of the C565T mutant (in the *rpoSp-kan* reporter). Data points represent measures of the frequency of the C565T mutant in a single culture. C565T mutants in the *rpoSp-kan* culture were confirmed via sanger sequencing. Black dots represent the mean values of 12 biological replicates (grey dots) and error bars one standard deviation from the mean**. B) Deletion of *psrA* reduces transcription from *rpoSp*.** SBW25 and SBW25 Δ*psrA* were grown as per fluctuation assays and RT-qPCR was performed using methods described in Fig. 3. SBW25 Δ*psrA* had ∼4 fold reduction in promoter activity from *rpoSp*. Black dots represent the mean values of six biological replicates (gray dots) and error bars one standard deviation from the mean.

### The high mutation rate is conditional on transcription from the *rpoSp*

Fluctuation assays identified high rates of the C565T mutation associated with *rpoSp*. Unknown is the underlying mechanism, and specifically whether transcription from *rpoSp* is the cause of elevated high mutation rate at base pair 565. We had identified the likely presence of *rpoSp* medial to *nlpD* using qPCR, and confirmed an increase in transcription from this region in stationary phase cultures (Fig. 3B), but lacking was an understanding of the spatial and functional relationship between C565T and *rpoSp*.

To this end, we mapped the transcription start site (TSS) of *rpoSp*. Using 5’RACE, the beginning of the transcript from *rpoSp* (in both SBW25 and SBW25 Δ*wss* was mapped to approximately 10 bp downstream of the C565T mutation (see Fig. S3 and Table S2). Transcription of *rpoSp* was previously mapped to this identical location in *P. aeruginosa* (Fujita et al., 1994). Similar to *P. aeruginosa*, *rpoSp* has no obvious −10 promoter site, but does share sequence similarity to the “gearbox” promoters - promoters induced by RpoD during periods of slow growth (Bohannon et al., 1991). Relevant for this study, the C565T mutation is predicted to alter the “gearbox” region, potentially altering transcription from *rpoSp*. Furthermore, the spatial relationship of C565T to the TSS holds strong similarity to a previously described study (Sankar et al., 2016), that identified high rates of T to C point mutations 7 bp outside the transcription start site of the transcriptionally induced gene *thyP3*.

Observing the C565T mutation affected the −10 region of *rpoSp* and resulted in higher transcription, the transcriptional effects of the Q189W (sequence TGG) mutation on transcription from *rpoSp* was measured by RT-qPCR. The Q189W mutation was associated with a reduced mutational parallelism (Fig. 1B). No measurable induction of expression from *rpoSp* was detected for cultures of SBW25 *Δwss nlpD* Q189W (Fig. S4). This Q189W mutation also further removed one of the consensus nucleotides comprising a “gearbox” promoter (Bohannon et al., 1991). These results provide further evidence the - 10 region of *rpoSp* as crucial for determining transcription from *rpoSp*, and supported the hypothesis that transcription from *rpoSp* caused the high C565T mutation rate.

A second region of *rpoSp* important for determining levels of transcription, and possibly rates of the C565T mutation, is the PsrA binding region. In *Pseudomonas*, transcription of *rpoS* is positively induced by the transcriptional regulator PsrA, which binds upstream of *rpoSp* and within *nlpD* (Kojic et al., 2002). The role of PsrA as an inducer of transcription from *rpoSp* was confirmed by quantitative RT-qPCR. Transcript from *rpoSp* was measured for cultures of SBW25 and SBW25 Δ*psrA* (Fig. 4B). The absence of PsrA reduced the transcription from *rpoSp* by ∼6-fold, demonstrating that the level of transcription from *rpoSp* is conditional on PsrA.

Given existence of the *rpoSp* within *nlpD*, positive activation by PsrA, and position of base pair 565, and the general association between mutational parallelism and transcription from *rpoSp*, we reasoned that transcriptional activity at the rpoS promoter was necessary for hyper-mutation at base pair 565. This hypothesis also drew on the finding of Sankar and colleagues, that induction of transcription could elevate rates of a particular T to C mutation −7 bp to the TSS of *thyP3* (Sankar et al., 2016). To this end, we inserted the ‘*rpoSp-kan’* mutational reporter into the SBW25 Δ*psrA*, and performed fluctuation assays identical to those with SBW25. The frequency of C565T mutants in culture was two orders of magnitude lower when PsrA was not present to induce transcription from *rpoSp* (Fig. 4A) right). The mutation rate of the C565T mutation in the background of SBW25 Δ*psrA* was 7.2 × 10^−9^ per base-pair per replication (95% CI = 9.2 – 5.4 × 10^−9^), as derived by a MSS Maximum Likelihood Method. This represents a 60-fold decrease compared to the rate of mutation in SBW25. As before, the fitness of these mutations was assess using relative fitness assays (Fig. S2). This revealed no advantage of the C565T mutations evolved in the SBW25 Δ*psrA* mutational reporter, suggesting mutation alone was responsible for the frequency of these mutants in fluctuation assays. Together, these measures demonstrate the high C565T mutation rate is conditional on *rpoSp* and the induction of transcription by PsrA.

## Discussion

The work described here shows the open reading frame of *nlpD* features a nucleotide at position 565 (codon 189) that mutates from cytosine to thymine at a rate 5,700-fold above the average per base-pair mutation rate for *P. fluorescens* SBW25 (Long et al., 2018). This high mutation rate was initially suspected following observations of near-deterministic levels of parallel evolution caused by the C565T mutation. Analysis of the fitness effects of a range of *nlpD* mutants showed that elevated fitness benefits due to the C565T mutation are insufficient to explain its frequent occurrence. Genetic analysis showed C565T occurs within the promoter sequence of the stationary phase sigma factor RpoS. The C565T mutation has two phenotypic effects: cells chaining arising from inactivation of *nlpD* and the elevation of transcription from *rpoSp*. This elevation in transcription allowed construction of a mutational reporter permitting measures of the C565T mutation rate. Finally, we show that this extreme mutation rate is dependent in the induction of transcription by the positive inducer PsrA.

The C565T mutation is remarkable both in terms of the specific location and rate. Instances of parallel evolution caused by elevated mutation rates are both predicted by theory (McCandlish & Stoltzfus, 2014) and are supported by numerous experimental studies (see references within (Horton & Taylor, 2023)). However, rarely are hotspots associated with a selectable phenotypic change that allow determination of specific substitutions rates (Barnett, Zeller, & Rainey, 2024; Kapel, Caballero, & MacLean, 2022; Sankar et al., 2016; Viswanathan, Lacirignola, Hurley, & Lovett, 2000). In the case of C565T, an estimate of the mutation rate was made possible following the fortuitous identification that the mutation altered transcription from *rpoSp*, and that this mutation rate was shown to be dependent on the presence of PsrA. We are not aware of reports of a mutational hotspot occurring within a wild-type promoter. Furthermore, this hotspot is remarkable in that is found within a region – the *nlpD*-*rpoS* pseudo-operon – which is highly conserved across gamma-proteobacteria (Gottesman, 2019).

The single study that bears greatest similarity to the results reported here is that by Sankar et al (Sankar et al., 2016). The authors, using the model species *Bacillus subtilis*, reported thymine to cytosine mutations occurring at a rate of ∼2.7 × 10^−8^ per cell per generation, specifically −7 to the transcription start site within the IPTG-inducible promoter of *thyP3*. In comparison, we identified cytosine to thymine mutations at a rate of ∼ 4.2 × 10^−7^ per base-pair per replication, approximately −10 to the transcription start site in a positively induced promoter. These positional similarities suggest a similar mutational process – Sankar and colleagues suggested solvents in contact with the template strand in the transcriptional bubble might initiate nucleotide lesions. In our case, mutagenesis is influenced by PsrA, which is required for the high C565T mutation rate. PsrA is predicted to bind to the PsrA binding site approximately (50 to 29 bp upstream of the C565T mutation), as informed by DNAse footprint assays in *P. putida* which share an almost identical binding site sequence (Kojic et al., 2002). Consequently, it is possible that the presence of PsrA may introduce a secondary effect and amplify mutation rates – such as blocking the mis-match repair process by MutS or MutL. We hypothesize that a systematic analysis of mutation rates in promoters with different regulation structures – possibly by taking advantage of techniques such as maximum-depth sequencing (Jee et al., 2016) – may reveal differential effects of promoter sequence variation on mutation rates.

Assuming that positive regulation of promoters may lead to higher local mutation rates, might this process speed up adaptive evolution? Assuming mutational hotspots are identifiable at other inducible promoters, these mutations may play a unique role in providing mutational fuel for adaptation. The spatially specific nature of these hotspots limits mutations to regions critical for transcription. It is possible that these hotspots might be especially active when promoters are induced – meaning that the hotspots are likely active in promoters which drive environmentally-relevant genes. This suggests a scenario in which bacteria, sensing environmental signals, might have high rates of mutation in promoters, resulting in variation to niche-relevant transcript levels. Such mutants stand to be presented to selection at a rate far higher than functional changes in open reading frames. Resulting promoter mutations might also revert at high rates under scenarios in which high transcript levels are no longer required. Such a scenario would explain experimental findings where promoter mutations are found at a high frequency in experimental populations (Lind, Farr, & Rainey, 2015; Pentz & Lind, 2021).

With focus on reporting high mutation rates within *rpoSp*, we have made little of the fact that this high mutation rate may have multiple ecological consequences in wild populations of *P. fluorescens* – or indeed in other gamma-proteobacteria – where the sequence of *rpoSp* is highly conserved. The C565T mutation has two functional consequences – cell chaining and the higher expression of *rpoS* transcript. While translational and post-translational regulation of RpoS might limit any impact of higher transcript levels (Gottesman, 2019), the truncation of NlpD has clear phenotypic effects. In our experimental system, chains of cells allow adaptation to static microcosms of rich media (Lind et al., 2017), but unknown is the fitness consequence of these chains in natural settings. Cell chaining is likely to be deleterious. Assuming the cells of each chain share a common periplasmic space, infection and lysis of any one cell by bacteriophage may result in the death of the entire chain. The presence of a mutational hotspot inside *rpoSp,* which likely has deleterious consequences on cellular morphology, seems antithesis to the highly conserved nesting of *rpoSp* within *nlpD* across gamma-proteobacteria (Gottesman, 2019). Assuming this hotspot is found in more gamma-proteobacteria, it would suggest a strong selective benefits for nesting *rpoSp* within *nlpD* – a benefit yet to be determined. We consider the nesting of *rpoSp* within *nlpD* – and the mutational hotspot there in – a profound mystery requiring understanding of the maintenance of complex genetic regulation over vast evolutionary time scales.

## Methods

### Media and strains

Derivatives of *Pseudomonas fluorescens* SBW25 were used in all experiments (Silby et al., 2009). The SBW25 Δ*wss* strain is identical to that previously described (Lind et al., 2017). *Escherichia coli* strains were used for the alteration of the SBW25 genome - *E. coli* Top10, *E. coli* DH5-α λ*_pir,_ E. coli* pRK2013, *E. coli* S17-1 λ*_pir_*. All strains were stored in ∼25% glycerol saline solution at −80 °C. Unless otherwise stated, *P. fluorescens* was cultured using King’s B (KB) medium (King, Ward, & Raney, 1954) and *E. coli* using lysogeny broth (LB) medium (Bertani, 1951). Media was solidified using 1.5% agar. Unless otherwise stated, liquid cultures were incubated with orbital shaking (220 rpm) at either 28 °C for ∼48 h (for *P. fluorescens*) or 37 °C for 24 h (for *E. coli*). Co-cultures of *E. coli* and *P. fluorescens* were incubated at 28 °C. Media was supplemented with the following selection agents for the construction of strains: kanamycin (50 mg l^−1^), nitrofurantoin (1 g l^−1^), tetracycline (15 mg l^−1^). Kanamycin was added to LB agar at 400 mg l^−1^ for selection of C565T mutants using the ‘*rpoSp-kan*’ cassette (see below). Double recombinants from two-step allelic exchange using our in-house pSACB plasmid were selected in TSY10 media (Huang & Wilks, 2017).

### Reconstruction of mutations and insertion of genes

The introduction of plasmids into *nlpD* loss-of-function strains failed after multiple attempts – it is important to note that complementation or transfer of similar *nlpD* mutations require transfer into another genotypic background to construct desired constructs.

To enable fitness assays, derivatives of *P. fluorescens* SBW25 were marked with *sgfp2* and *mScarlet-I* at the neutral *attTn7* locus (Choi & Schweizer, 2006) using plasmids described in detail elsewhere (Schlechter et al., 2018). *E. coli* S17-1 λ*_pir_* hosting plasmids pMRE-Tn7-152 or pMRE-Tn7-155 were cultured overnight with selective antibiotics along with target SBW25 genotypes. Bi-parental mating was then performed on the resulting cultures (see below). Transconjugants of *P. fluorescens* were selected on LB agar containing 0.1 % W/V arabinose, nitrofurantoin and kanamycin to select for transposition of the mobile section of pMRE-Tn7-152 or pMRE-Tn7-155 into the *attTn7* site. Resulting fluorescent colonies were then streaked on LB agar containing kanamycin, and resulting colonies were assessed by PCR for integration at the *attTn7* site. Overnights from these colonies were made with selective kanamycin and stored at −80 °C.

Reconstruction of mutations, or the deletion of genes, was achieved using two-step allelic exchange using sucrose counter-selection (Hmelo et al., 2015). To reconstruct existing mutations in other genotypes, template containing the mutation of interest was amplified with primers approximately 500 - 800 up and downstream of the mutation targeted for reconstruction (i.e. approximately 1400 bp in total) using Q5 high fidelity taq polymerase (NEB). In the case of reconstruction of *nlpD* mutations evolved from Q189W, the Q189W mutation was reverted to the WT sequence by strand overlap extension PCR (Ho, Hunt, Horton, Pullen, & Pease, 1989). To delete genes, similar sized regions either size of the gene of interest were amplified, using primers with complementary tails at one end of each fragment. These fragments were cloned using an inhouse plasmid ‘pUIsacB’ (GenBank accession number pending), which features genes for tetracycline resistance (*tetA*), fluorescence (*msfgfp*) and sucrose counter selection (*sacB*). pUIsacB was linearized using restriction enzymes and then amplified using Q5 polymerase. Amplified pUIsacB was digested with DpnI. The *nlpD* fragments were amplified with primers with a 5’ tail complementary to 20 nucleotides to the ends of the pUIsacB amplicon. Both amplicons were purified using a ‘Qiaquick PCR purification kit’ (Qiagen), product concentrations were quantified, and then joined into a circularized plasmid by ‘NEBuilder HiFi DNA assembly’ (NEB) following manufacturers protocols. Assembled products were transformed into chemically competent *E*. *coli* Top10 cells, with transformed cells plated on LB agar containing tetracycline. Several colonies from each transformation were subject to colony PCR to confirm the correct sized insert and several colonies were used to inoculate overnight cultures. Overnight cultures were also started of *E. coli* pRK2013 (using LB supplemented with kanamycin), and the target host strain of the reconstruction (SBW25 Δ*wss*). Resulting *E. coli* pUIsacB*-nlpD* cultures were then frozen, and the pUIsacB-*nlpD* was transferred to the target host by tri-parental mating (see below), with transconjugants selected with LB agar plates supplemented with tetracycline and nitrofurantoin. Several resulting colonies were streaked on TSY10 plates to counterselect for pUIsacB recombinants. Resulting colonies were scanned for the phenotypic effects of the reconstructed mutation – in this case cell chaining identified by microscopy – and colonies were streaked on KB agar plates, along with LB-tet plates to confirm loss of pUIsacB. Colonies of the host strain with the recombined mutation were inoculated into KB microcosm for storage. PCR and sanger sequencing was performed on all reconstructed mutants to ensure the correct mutation (and the absence of additional mutations) at the intended locus.

Sucrose counter selection was not used for construction of SBW25 Δ*wss nlpD* C565T A566G, or for reconstruction of *nlpD* mutations into SBW25 Δ*wss* (and as template for subsequent reconstruction using‘pUIsacB’) which was made using a previously described two-step allelic exchange method (Bantinaki et al., 2007; Rainey, 1999). Briefly, forward and reverse primers with binding sites ∼800 bp from the C565T A566G mutation were used to amplify the template DNA. This fragment was ligated into the tetracycline-resistant pUIC3 plasmid (Rainey, 1999) and cloned in *E. coli* DH5-α λ*_pir_*. Conjugation with *E. coli* pRK2013 was used to transfer the pUIC3 construct into SBW25 Δ*wss*, and single recombinants were selected for with tetracycline and nitrofurantoin. Recombinant clones were used to initiate cultures which were treated with cycloserine with counter-selected single recombinants. The culture was plated and single colonies were stored which were cured of the pUIC3 plasmid, and the absence of mutational scars and the presence of the desired mutation in *nlpD* was confirmed by sanger sequencing.

Construction of the ‘*rpoSp-kan* reporter’ and fluorescent strains with this reporter were also constructed using ‘NEBuilder HiFi DNA assembly’ (NEB). To construct the ‘*rpoSp-kan* reporter’, 401 bp of *nlpD* (with the C565T mutation situated centrally) was amplified along with a kanamycin resistance gene (kanR) and a backbone plasmid (extracted from MPB15151, see (Theodosiou, Farr, & Rainey, 2023)) encoding Tn7 elements, an origin of replication and a tetracycline resistance enzyme. These three fragments were assembled using ‘NEBuilder HiFi DNA assembly Master Mix’ (NEB) (see above), and transformed into top10 chemically competent cells. The resulting plasmid (*pTn7-rpoSp-kan*, genbank number pending) was extracted from liquid couture via miniprep using a QIAprep Spin Miniprep Kit (Qiagen) transformed along with the pUX-BF13 (to express transposition machinery, (Bao, Lies, Fu, & Roberts, 1991) into derivatives of SBW25 using electroporation (protocol previously described (Choi & Schweizer, 2006). Resulting transformants were selected using LB agar supplemented with tetracycline and the insertion of the construct at the *attTn7* was confirmed by PCR and electrophoresis. The integrated *rpoSp-kan* genetic construct validated with sanger sequencing. The Tn7-*rpoSp-kan* reporter plasmid was similarly modified to express either *sGFP2* or *mScarlet-I* (resulting in plasmids *pTn7-gfp-rpoSp-kan*, or *pTN7-scarlet-rpoSp-kan,* genbank numbers pending), and this construct was again integrated into the *attTn7* region of SBW25 to enable fitness assays. Attempts to introduce a C565T mutation into *pTn7-rpoSp-kan* failed in multiple attempts, and spontaneous evolution was used to derive *nlpDC565T-kan, gfp-nlpDC565T-kan* and *pTN7-scarlet-nlpDC565T-kan*.

Transfer of plasmids from *E. coli* cloning strains to derivatives of *P. fluorescens* SBW25 was performed with tri-parental or bi-parental mating or by electroporation where specified. Triparental mating required overnight cultures of the donor strain with the intended plasmid, a helper strain (*E. coli* pRK2013) and the target *Pseudomonas* host strain. Where the donor was *E. coli* S17-1 λ*_pir_*, the helper strain was omitted. When the cultures were grown to saturation, 1mL samples of SBW25 culture was heat shocked at 45 °C for 20 mins. Meanwhile, 500 µL of *E. coli* culture was washed of remaining selective antibiotics by centrifugation, and resuspension of the cells in 500 µL of LB. The heat-shocked SBW25 culture was then added to the *E. coli* culture, and the mixed culture was centrifuged, supernatant was removed and the cells resuspended in 100 µL of LB. The resuspension of cells was then spread over a ∼4 cm diameter region of a plate of LB agar and incubated for ∼24 h. Transconjugants were then resuspended in 1 mL of LB and dilutions were plated on selective LB agar plates as indicated below to select for transconjugants.

### Experimental evolution of the C565T mutants

The suite of *nlpD* mutants evolved from SBW25 Δ*wss* and SBW25 Δ*wss nlpD* C565T A566G were evolved using previously described methods (see (Lind et al., 2017)). Extra care was taken to limit contamination with a potential C565T mutation in the SBW25 Δ*wss* inocula. Accordingly, individual colonies of SBW25 Δ*wss* were used to inoculate individual KB microcosms, which were then incubated statically for five days. The genotype SBW25 Δ*wss nlpD* C565T A566G was inoculated direct from an isogenic frozen stock into replicate microcosms of KB media. These microcosms were incubated for six days and a sample of the microcosm was stored at −80 °C, and dilutions of the microcosms were plated over several KB plates. Resulting colonies were assessed by eye for ancestral appearing morphologies with subtle alteration in opaqueness, and different morphotypes were assessed by microscopy for a chaining cellular phenotype. Individual mutant colonies were then streaked on KB agar plates to produce single colonies which were used to produce cultures from which *nlpD* was sanger sequenced.

### Fitness assay of *nlpD* mutants

Fitness assays of *nlpD* mutants – or *rpoSp-kan* constructs – were performed as previously described (Lind et al., 2015), with the following exceptions. For fitness assays of the ‘*rpoSp-kan’* mutants, cultures were prepared to replicate fluctuation assays, with 24 hr incubated shaking cultures of each competitor mixed (50:50), sampled for flow cytometry and plated on KB agar, and the competitors diluted 10^−7^ in new KB microcosms to found the test microcosms. These test microcosms were then sampled again (with flow cytometry and plating on KB agar) after 22 h of shaking at 220rpm. Fitness assays compared strains marked with sGFP2 or M-Scarlet-I (Schlechter et al., 2018)), with fluorescent proteins swapped for half the replicates to correct for fitness cost of the fluorophores. Fitness assays in statically incubated microcosms also included an equal ratio of non-marked ancestral to help recreate the conditions in which *nlpD* mutants evolved. The relative frequency of either fluorescent marked mutant was measured using a ‘MACSQuant VYB’ flow cytometer (Miltenyi). Samples were run with a ‘low’ flow rate, ‘moderate’ mixing and ‘strandard’ cleaning between samples. Samples were prepared for flow cytometery diluted with filtered PBS buffer in 96-well plate wells in order to achieve ∼500-800 recorded events per second. The cytometer had a trigger set at value 1.4 of the SSC channel, and events were gated to not include events from the filtered PBS buffer used to dilute samples, and at least 20,000 events were sampled. Events were plotted of events from 525/50 nm (sGFP2) and 615/20 nm (M-Scarlet-I) detectors. We calculated selection coefficients per generation with the regression model s = [ln(R(t)/R(0))]/[t], in which R is the ratio of each compositor at tested time points, and t is the number of generations (Dykhuizen, 1990). Fitness assays were performed with 8 replicates performed over 2 separate occasions.

### Rapid amplification of cDNA ends

Triplicate cultures of SBW25 and SBW25 Δ*wss* were each inoculated with a single colony and incubated statically for 24 h. Samples of SBW25 and SBW25 Δ*wss* (200 µL and 400 µL respectively) were taken, and total RNA was extracted following manufacturer’s instructions using an RNeasy Mini Kit (Qiagen) in conjunction with RNAprotect tissue reagent (Qiagen). The 5’ end of *rpoS* mRNA was then mapped using a ‘5’ RACE System for Rapid Amplification of cDNA, version 2.0’ (Invitrogen). A sample of 1 µg of total RNA as the sample input, with the first extension primer directly outside of the *nlpD* open reading frame. PCR products were amplified over two rounds, and resulting cDNA was amplified with *taq* polymerase (NEB) to allow cloning of the resulting products into pCR2.1 using a ‘TOPO TA Cloning Kit’ (Invitrogen), with plasmids transformed into top10 chemically competent cells. At least 10 resulting colonies were stored and colony PCR was used to amplify the region of insert and an insert-annealing primer used for a sanger sequencing reaction to identify the 5’ start of the mRNA transcript.

### Quantitative RT-PCR

Quantification of transcript levels across *nlpD* was performed using qPCR using previously described methods (Lind et al., 2015). Briefly, total RNA was isolated from shaken cultures grown either to specified time points or to stationary phase (OD_600_ = 2.5 to 3.0). RNA was isolated using the SV Total RNA isolation system (Promega). Cultures were centrifuged, and pellets were resuspended in TE buffer (10 mM Tris, 1 mM EDTA pH 8.0) with 0.4 mg mL^−1^ lysozyme. The recommended protocol was then performed for extracting RNA from gram negative bacteria. Total RNA was then reverse transcribed using to DNA using a ‘High capacity cDNA reverse transcription kit’ (Applied biosystems). Resulting cDNA was then diluted 1:40 before use as template in the qPCR reaction (8 µL of diluted template per 20 µL reaction) using ‘PowerUp SYBR Green Master Mix’ (Applied biosystems). Reactions were performed in “MicroAmp Optical 96-Well Reaction Plates” (Applied biosystems) and reactions were measured in a “Quantstudio 1 Real-Time PCR System” (Applied biosystems). The change in transcript downstream of *rpoSp* was made relative to transcript upstream of *rpoS*, but within *nlpD*, using the ΔΔCq method (Livak & Schmittgen, 2001). Two replicate qPCR measures were made of each biological replicate, with each run performed on a separate occasion.

### Fluctuation assays

In order to measure the rate of mutation of the C565T mutation, a construct (*rpoSp-kan*, see methods above) was employed which would allow selection of the mutation in *rpoSp* while removing *rpoSp* from the fitness-altering context of *nlpD*. Six individual transformants of the *rpoSp-kan* construct in both SBW25 and SBW25 Δ*psrA* were used for fluctuation assays, each used on two separate occasions. For each replicate measure, individual colonies of the reporter strain were grown (without antibiotic selection) and then inoculated into shaking KB cultures for 24 h. Cultures were diluted 1 × 10^−7^ into 6 mL microcosms of KB media, and this culture was immediately plated to derive estimates of the inoculum (typically ∼1000 CFU mL^−1^). Cultures were then shaken for 22h, and then diluted by 1 × 10^6^ and plated on KB plates to get measures of CFU. Cultures of SBW25 were diluted 100 and 1000-fold for selective plating on multiple LB plates with kanamycin (400 mg l^−1^), while SBW25 Δ*psrA* mutants were plated on selective plates without dilution. Duplicate controls genotypes (cultured the same as the above) which already feature the C565T mutants were also plated with a 1 × 10^−6^ or 1 × 10^−7^ dilution of selective plates. All plates were incubated for ∼72 h at 28 °C. The total number of cells per plated culture were estimated from CFUs on KB media, while selective plates were visually scanned for colonies with a similar size to the control plates. To minimise the possibility of spontaneous mutants arising during slow growth on the selective plates, a minimum size of colony was selected as candidate colonies with C565T mutations in the reporter (3 mm or 4 mm was the minimum diameter size on plates with respectively more or less than 100 visible colonies). Candidate colonies were the marked and counted, and a random selection of 8 such colonies were selected for sanger sequencing. Upon identification of the C565T mutation in the construct, the total number of candidate colonies was corrected by the fraction of confirmed C565T mutants to produce an estimate of the true number of C565T mutants in the population. MSS maximum likelihood methods were used to analyse mutation rates using ‘FALCOR’ (https://lianglab.brocku.ca/FALCOR/; (Hall, Ma, Liang, & Singh, 2009)).

### Data accessibility

Data analysis was performed in ‘Rstudio’ (Posit team, 2023) version 2023.9.1.494, utilising ‘R’ (R Core Team, 2023) version 3.1.2, and data visualisation was performed using packages ‘ggplot2’ (Wickham, 2016). Raw data and calculations performed to produced visualised data are available via Zenodo (Farr, 2024), with each figure corresponding to a different folder. Sequences of the pUIsacB and ‘*rpoSp*-*kan’* selectable reporter will be deposited in GenBank upon peer-review.

## Acknowledgements

Thanks to David Rogers and Ellen McConnell for advice during molecular cloning, Elisa Brambilla for laboratory management and the MPI for Evolutionary Biology sequencing team. ADF, CV and PBR acknowledge generous core funding from the Max Planck Society.

