## Supplementary Figures and Tables for "An extreme mutational hotspot in *nlpD* depends on transcriptional induction of *rpoS*"

11.07.2024

[1]Department of Microbial Population Biology, Max Planck Institute for Evolutionary Biology, Plön 24306, Germany.

[2]New Zealand Institute for Advanced Study, Massey University, Auckland 0745, New Zealand

[3]Department of Molecular Biology, Umeå University, 901 87 Umeå, Sweden

[4]Umeå Centre for Microbial Research (UCMR), Umeå University, 901 87 Umeå 901 87, Sweden

[5]Laboratoire Biophysique et Évolution, CBI, ESPCI Paris, Université PSL, CNRS, 75005 Paris, France.

*Corresponding authour:

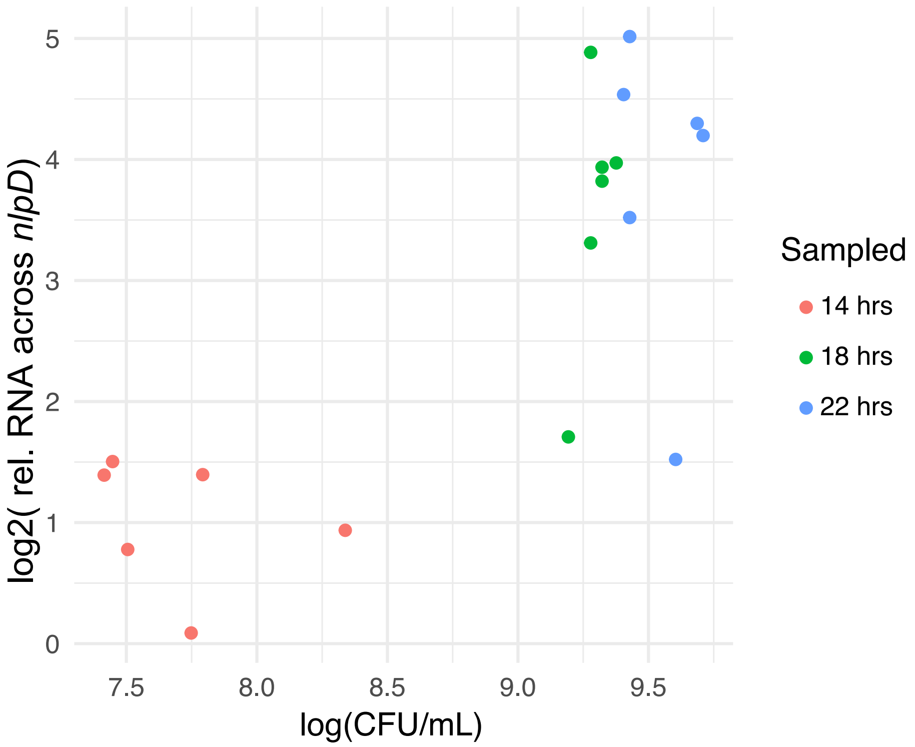


**Figure S1: Growth of SBW25 causes induction of transcription from *rpoSp*.** SBW25 was grown from a small inoculum (~1000 CFU $mL^{-1}$) for 14, 18 and 22 h and samples of the cultures were used to measure CFU concentrations, and also processed (as per Fig. 3) to measure induction of transcription from *rpoSp* relative to a region in *nlpD* upstream of *rpoSp*. Similar levels of activity from *rpoSp* were measured after 18 and 22 hours of growth, and fluctuation assays used cultures grown for 22 hours to ensure cells were in stationary phase.

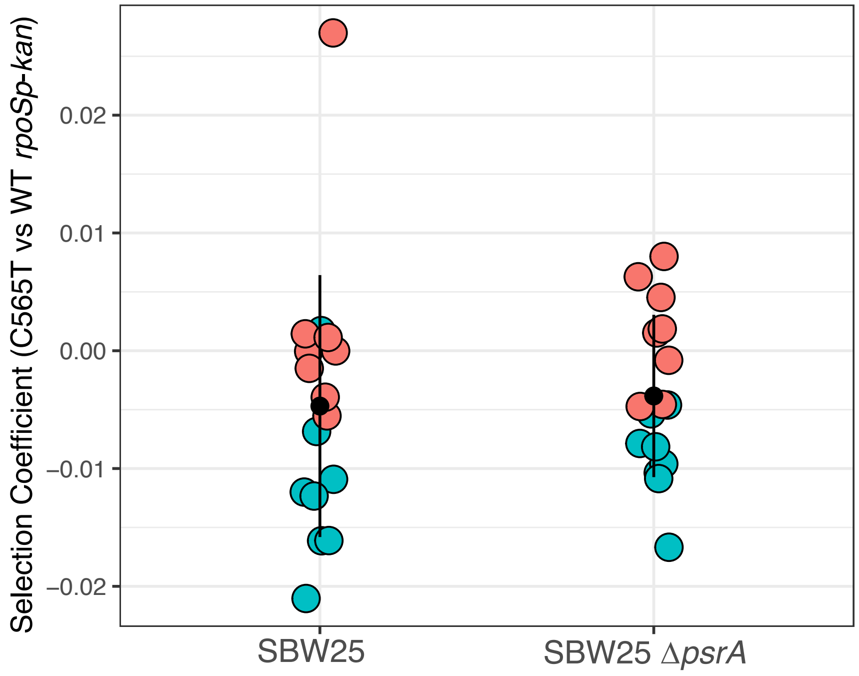


**Figure S2: Relative fitness of SBW25 genotypes with C565T mutated *rpoSp-kan* reporters vs non-mutant reporter genotypes.** The *rpoSp-kan* reporter construct was made to express Green or mScarlet fluorescent proteins, and evolved in genotypes SBW25 or SBW25 *ΔpsrA* to feature a C565T mutation in the *rpoSp-kan* reporter. Presented is the relative fitness of genotypes with mutant vs unmutated *rpoSp-kan*. Competitions were initiated with a 1:1 ratio and grown in a manner similar to fluctuation assays. Reciprocal pairwise competitions were used, with the C565T mutation in either a GFP (green) or mScarlet (red) background. A significant lower fitness was measured for the C565T mutant in *rpoSp-kan* in both backgrounds (Wilcoxon signed rank exact test of either SBW25 or SBW25 *ΔpsrA*, V = 29, p-value = 0.04431). Each competition involved 16 replicates, black dots represent the mean values and error bars one standard deviation from the mean.

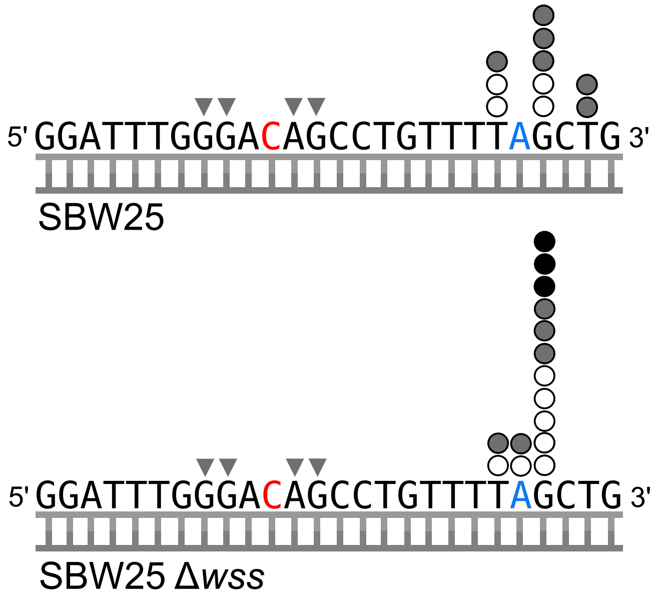


**Figure S3. The position of the C565T mutation relative to the transcriptional start site (TSS).** The TSS of *rpoS* is approximately 12bp downstream of the position of the C565T mutation (Red C). Circles represent the TSS from individually cloned reverse-transcription fragments from RNA expressed in stationary phase cultures of either SBW25 or SBW25 Δ*wss*. Cloned fragments were derived from one of three replicate cultures (indicated by either white, grey or black circles). The position of the mapped transcription start site in *P. aeruginosa* is marked in Blue (Fujita et al., 1994), triangles represent the consensus 'gearbox' promoter motif (Vicente, Kushner, Garrido, & Aldea, 1991).

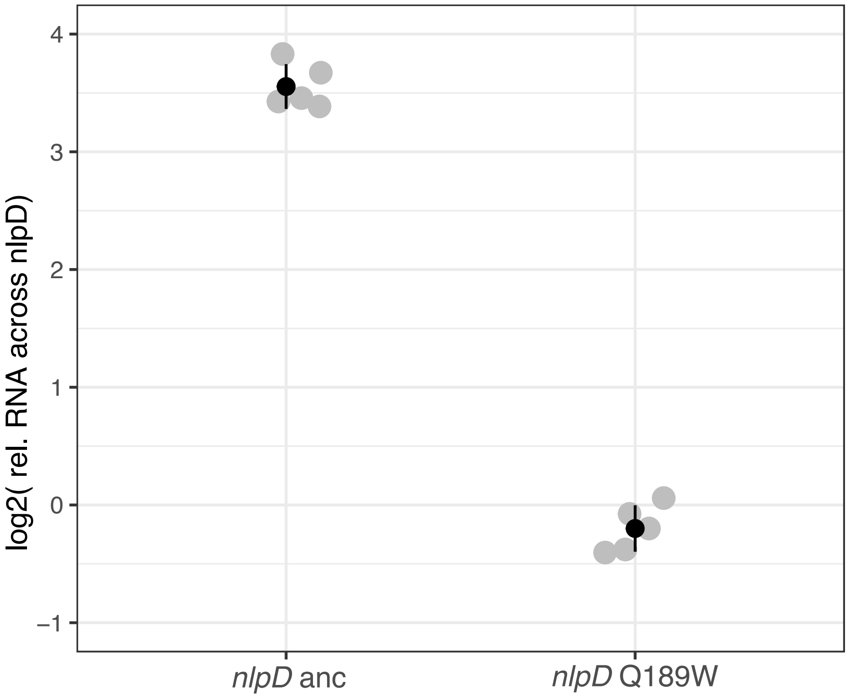


**Figure S4: The Q189W mutation prevents transcription from *rpoSp* for cells in stationary phase.** SBW25 Δ*wss* (*nlpD* anc) and SBW25 Δ*wss nlpD*Q189W (codon sequence TGG, which has an alteration of the *rpoSp* sequence) were grown to stationary phase, and mRNA was extracted, reverse transcribed and the regions up and down stream of *rpoSp* were used as templates for qPCR. SBW25 Δ*wss* has levels consistent with other measures of induced transcription from *rpoSp*, while the C565T A566G mutation to *rpoSp* removes any indication of transcription from *rpoSp* (i.e. there is no difference in the levels of transcript downstream compared to upstream of *rpoSp*). Black dots represent the mean values of five biological replicates (gray dots) and error bars one standard deviation from the mean.

**Table S1. Mutational positions and residue changes of *nlpD* mutants arising from populations of SBW25 ∆*wss nlpD* (Q189W) grown in statically incubated microcosms.** *indicates mutations reconstructed for fitness assays. Numbers within brackets represents amino acids until a stop codon is reached.

| Isolate | Nucleotide change | Amino acid change |
|---|---|---|
| 1 | G553A | G185R |
| 2* | Δ487-570 | Δ163-190 (Δ 28aa) |
| 3* | C603G | Y201* |
| 4, 14, 27 | A680G | H227R |
| 5, 6, 7, 9, 13, 16, 20, 22 | G566A | Q189* |
| 8 | Δ623-634 | Δ208-212 (Δ ATAS) |
| 10 | C775G | H259N |
| 11* | G483A | W161* |
| 12 | Δ620 | ΔR207(146) |
| 15 | Δ462 | P154(13) |
| 17 | G677A | G226D |
| 18 | C212T | A71V |
| 19 | C787T | R263C |
| 21, 26, 31 | A602G | Y201C |
| 23 | C811T | P271S |
| 24* | C817T | Q273* |
| 25, 30 | G539A | G180D |
| 28 | - | - |
| 29 | G493A | G165R |
| 32 | T91C | C31R |
| 33* | Δ396-397 | P132(43) |
| 34 | Δ141-144 | A47(6) |
| 35 | C484T | P162S |
| 36 | G595T | V199L |
| 37* | C49T | R17* |
| 38 | A809G | D270G |

**Table S2 : Positioning of the transcriptional start site (TSS) from total RNA extracts of SBW25 and SBW25 Δ*wss* using 5-prime RACE.**

| Genotype | MPB number of isolate | Replicate | Colony | Position in *nlpD* ORF |
|---|---|---|---|---|
| SBW25 | MPB29692 | 1 | 1 | 666 |
| SBW25 | MPB29693 | 1 | 2 | 577 |
| SBW25 | MPB29694 | 1 | 3 | 591 |
| SBW25 | MPB29695 | 1 | 4 | 575 |
| SBW25 | MPB29696 | 1 | 5 | 644 |
| SBW25 | MPB29697 | 1 | 6 | 744 |
| SBW25 | MPB29698 | 1 | 7 | 575 |
| SBW25 | MPB29699 | 1 | 8 | 659 |
| SBW25 | MPB29700 | 1 | 9 | 648 |
| SBW25 | MPB29701 | 1 | 10 | 577 |
| SBW25 | MPB30639 | 2 | 1 | 577 |
| SBW25 | MPB30640 | 2 | 2 | 577 |
| SBW25 | MPB30641 | 2 | 3 | 575 |
| SBW25 | MPB30642 | 2 | 4 | 708 |
| SBW25 | MPB30643 | 2 | 5 | 659 |
| SBW25 | MPB30644 | 2 | 6 | 664 |
| SBW25 | MPB30645 | 2 | 7 | 579 |
| SBW25 | MPB30646 | 2 | 8 | 577 |
| SBW25 | MPB30647 | 2 | 9 | 698 |
| SBW25 | MPB30648 | 2 | 10 | 579 |
| SBW25 | MPB30649 | 3 | 1 | 696 |
| SBW25 | MPB30650 | 3 | 2 | 734 |
| SBW25 | MPB30651 | 3 | 3 | No product |
| SBW25 | MPB30652 | 3 | 4 | 729 |
| SBW25 | MPB30653 | 3 | 5 | 500 |
| SBW25 | MPB30654 | 3 | 6 | 697 |
| SBW25 | MPB30655 | 3 | 7 | 714 |
| SBW25 | MPB30656 | 3 | 8 | 698 |
| SBW25 | MPB30657 | 3 | 9 | 422 |
| SBW25 | MPB30658 | 3 | 10 | 698 |
| SBW25 Δ*wss* | MPB29704 | 1 | 1 | 577 |
| SBW25 Δ*wss* | MPB29705 | 1 | 2 | 491 |
| SBW25 Δ*wss* | MPB29706 | 1 | 3 | 577 |
| SBW25 Δ*wss* | MPB29707 | 1 | 4 | 577 |
| SBW25 Δ*wss* | MPB29708 | 1 | 5 | 577 |
| SBW25 Δ*wss* | MPB29709 | 1 | 6 | 577 |
| SBW25 Δ*wss* | MPB29710 | 1 | 7 | 756 |
| SBW25 Δ*wss* | MPB29711 | 1 | 8 | 576 |
| SBW25 Δ*wss* | MPB29712 | 1 | 9 | 377 |
| SBW25 Δ*wss* | MPB29713 | 1 | 10 | 575 |
| SBW25 Δ*wss* | MPB30659 | 2 | 1 | 577 |
| SBW25 Δ*wss* | MPB30660 | 2 | 2 | 575 |
| SBW25 Δ*wss* | MPB30661 | 2 | 3 | 577 |
| SBW25 Δ*wss* | MPB30662 | 2 | 4 | 698 |
| SBW25 Δ*wss* | MPB30663 | 2 | 5 | 577 |
| SBW25 Δ*wss* | MPB30664 | 2 | 6 | 663 |

| | | | | |
|---|---|---|---|---|
| SBW25 Δ*wss* | MPB30665 | 2 | 7 | 648 |
| SBW25 Δ*wss* | MPB30666 | 2 | 8 | 577 |
| SBW25 Δ*wss* | MPB30667 | 2 | 9 | 576 |
| SBW25 Δ*wss* | MPB30668 | 2 | 10 | 698 |
| SBW25 Δ*wss* | MPB30669 | 3 | 1 | 711 |
| SBW25 Δ*wss* | MPB30670 | 3 | 2 | 643 |
| SBW25 Δ*wss* | MPB30671 | 3 | 3 | 698 |
| SBW25 Δ*wss* | MPB30672 | 3 | 4 | 577 |
| SBW25 Δ*wss* | MPB30673 | 3 | 5 | No product |
| SBW25 Δ*wss* | MPB30674 | 3 | 6 | 725 |
| SBW25 Δ*wss* | MPB30675 | 3 | 7 | 663 |
| SBW25 Δ*wss* | MPB30676 | 3 | 8 | 349 |
| SBW25 Δ*wss* | MPB30677 | 3 | 9 | 577 |
| SBW25 Δ*wss* | MPB30678 | 3 | 10 | 738 |
